# Semi-supervised Contrastive Learning for Bioactivity Prediction using Cell Painting Image Data

**DOI:** 10.1101/2024.10.26.620435

**Authors:** David Bushiri Pwesombo, Carsten Beese, Christopher Schmied, Han Sun

## Abstract

Morphological profiling has recently demonstrated remarkable potential for identifying the biological activities of small molecules. Alongside the fully supervised and self-supervised machine learning methods recently proposed for bioactivity prediction from Cell Painting image data, we introduce here a semi-supervised contrastive (SemiSupCon) learning approach. This approach combines the strengths of using biological annotations in supervised contrastive learning and leveraging large unannotated image datasets with self-supervised contrastive learning. SemiSupCon enhances downstream prediction performance of classifying MeSH pharmacological classifications from PubChem, as well as mode of action and biological target annotations from the Drug Repurposing Hub across two publicly available Cell Painting datasets. Notably, our approach has effectively predicted the biological activities of several unannotated compounds, and these findings were validated through literature searches. This demonstrates that our approach can potentially expedite the exploration of biological activity based on Cell Painting image data with minimal human intervention.

## 1 Introduction

The Cell Painting (CP) assay is a high-content screening method that enables an unbiased and high throughput evaluation of pharmacological similarity among compounds by comparing the drug-induced morphological changes on cells.^1^ The microscopy images generated by using this assay serve as the basis for image-based profiling, with the goal of capturing the rich information contained within the images and transforming them into multidimensional phenotypic profiles. Compounds with similar profiles are anticipated to induce similar perturbations of cell functions. Consequently, profiles of compounds with known mode of actions (MoAs) can be used to identify uncharacterized compounds that share related bioactivity.^2,3^ In contrast to traditional target-based screening approaches, profiling with transcriptomics or morphological image features can be used to provide a broad and unbiased systematic readout of the bioactivity of compounds, including information related to unknown targets or targets that pose challenges in target-specific assays. High-content morphological profiling has recently found numerous applications in identifying new bioactive compounds^4–6^, repurposing drugs^7–9^, and detecting toxic compounds^10–13^, promising to significantly expedite the drug discovery process.

Analyzing and exploring bioactivity from high-throughput microscopy images is generally challenging due to the high dimensionality of the image features and the complexity of the relationships among them. Traditionally, expert-engineered profiles extracted by software such as CellProfiler^14^ have been used for calculating representations of treatments in the Cell Painting assay. Recently, learning the profiles directly from images using machine learning (ML), and more specifically deep learning, has proven beneficial as it more effectively leverages image-based information.^2,7^ Additionally, ML classifiers can learn the relationship between the drug-induced perturbations and the bioactivity of annotated compounds, which can then be used to predict the bioactivity of unannotated compounds at scale (Figure 1). However, accurately predicting the bioactivity of compounds remains a significant challenge, as many compounds lack biological annotations or exhibit poly-pharmacology.^2,20^ Bioactivity prediction is further complicated by the fact that compound bioactivity can be reported at various levels of biology. Some annotations directly relate to the targets of the compounds, while others may indicate the pathways they modulate or more broadly the drug indications.^15^

**Figure 1.**
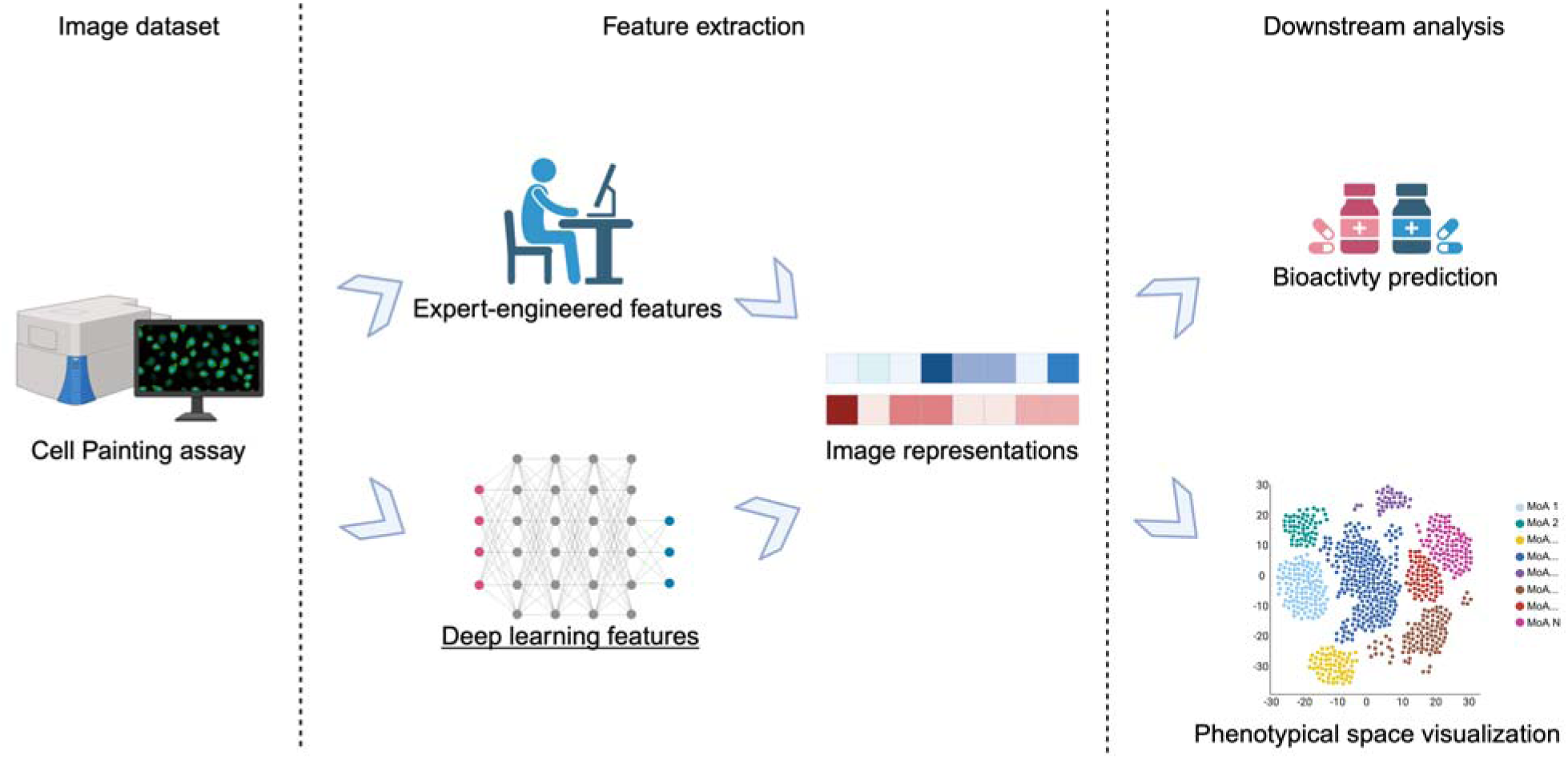
Two general workflows for learning and analyzing image representations, from Cell Painting microscopy images. (Top) Expert-engineered features calculated from software such as CellProfiler and (down) deep learning derived features. In the downstream analysis, the calculated representations can be used to visually analyze the relationship between the different compounds and to use them as features for a downstream machine learning model for predicting the bioactivity of the compounds.

Incorporating bioactivity labels during neural network training of the CP microscopy images can enhance the quality of the learned representations, as more information is provided to the neural network.^2^ Neural networks are often trained using a supervised approach, where they directly learn the relationship between the input data and the corresponding ground truth annotations. Several fully supervised ML approaches have recently been proposed to predict the bioactivity of compounds from CP image data.^21,22^ Supervised approaches generally achieve higher accuracies compared to other strategies, provided a large number of labels is available. However, supervised models can only be trained on annotated data points. This limitation poses a significant challenge for tasks such as bioactivity prediction, where labelled data is scarce.^2,23^

In contrast, unsupervised and self-supervised learning emerge as viable alternatives for training neural networks without the necessity of ground truth labels. This allows for the training of neural networks on considerably larger, unannotated datasets. In self-supervised learning, the neural network learns suitable representations of the input data by performing pretext-tasks. Such pretext tasks can be used to generate pseudo labels, by making use of additional metadata (e.g. cell type, perturbation)^2^ or by using the inherent structure of the data (e.g. augmenting images)^24^. Generating these pseudo-labels does not require expensive human annotation, thus enabling the creation of more data-efficient models.^24^ The learned representations are then typically used to train downstream models, which are then evaluated on their performance on specific tasks of interest. Recently, several self-supervised strategies have matched or even surpassed supervised models in various image analysis tasks, requiring only a fraction of the annotated data compared to supervised models.^24^ This underscores the effectiveness of self-supervised learning, particularly in tasks with limited annotations. In the context of applying representation learning for calculating image-based profiles, Perakis *et al*.^25^ introduced a self-supervised contrastive learning-based method. In contrastive learning, features from multiple images are simultaneously compared against each other, with the task of the neural network being to distinguish positive examples from negative ones. Positives can be generated by augmenting images or using different replicates of images.^26^ All other remaining images are defined as negatives. More recently, DINO^27^ (self-**di**stillation with **no** labels), a self-supervised algorithm has been shown in several studies to improve the downstream prediction of compound bioactivity using microscopy images from CP assays as input.^28–30^ Originally developed to learn representations of natural RGB images, DINO has achieved state-of-the-art performance in self-supervised methods across various computer vision tasks.^27,29^ One key difference between DINO and classical contrastive learning-based strategies is that DINO relies solely on pairs of positives to learn a similarity measure of images. As a result, it does not make use of negatives and does not simultaneously discriminate between multiple images.^27^ Although self-supervised methods have achieved state-of-the-art performance in many tasks, there are still certain cases where the predictive performance of models relies on training the neural network with labelled data.^31^ Classification in complex biological applications, such as bioactivity prediction from CP data, falls into this category.

To train a convolutional neural network (CNN) with limited labelled data, we introduce a novel approach in this study for learning image-based profiles using semi-supervised contrastive learning. This approach leverages a large pool of unlabeled images while also incorporating information from the labels of annotated images to train a convolutional neural network, thereby combining the advantages of both supervised and self-supervised methods. Semi-supervised methods have proven to be an effective strategy for image classification, as demonstrated by the Meta Pseudo Labels method by Pham *et al*., which achieved state-of-the-art performance for image classification on the ImageNet benchmark in 2021.^32^ Recently, semi-supervised strategies have also shown to be an efficient strategy in single-cell transcriptomics, for increasing performance on downstream classification tasks and inferring labels for unannotated data points.^33,34^ In this work, we took advantage of the multiple fields of views and replicates of treatments to define positive examples during contrastive learning. For supervised contrastive learning, we additionally consider pairs of data points as positives if they share the same label. We used Medical Subject Heading (MeSH)^35,36^ pharmacological classes from PubChem^37^ as labels for supervised contrastive learning.

A wide range of biological processes can be analyzed from Cell Painting image data, depending heavily on the specific research question. Currently, there are several prediction tasks and annotation systems used in the literature to evaluate bioactivity prediction from Cell Painting data, but there is no unified prediction task for evaluating machine learning models. This diversity in tasks and annotations reflects the complexity and breadth of potential applications, but it also highlights the challenge of developing standardized evaluation strategies for assessing model performance.

In this work, we focused on multi-label classification to better account for the polypharmacology of compounds. We performed downstream classification on three annotation systems (MeSH Classes, Drug Repurposing Hub MoA, and target annotations), each containing hundreds of different classes, and compared our approach with alternative methods for generating representations from Cell Painting image data. Our comparative analysis shows that our workflow improves downstream prediction across all evaluated tasks compared to using CellProfiler^14^, DINO^27^ (implemented comparable to Doran *et al.*^29^), and a contrastive learning-based approach^38^ as baselines. Notably, the DINO approach^29^ also used multi-label classification of Drug Repurposing Hub MoA annotations as downstream task. Additionally, we systematically assessed which MeSH classes could be predicted with sufficient precision using our workflow. Finally, we predicted pharmacological classes of unannotated compounds in the datasets. For the top-ranked compounds, we confirmed several predictions through literature searches, highlighting the effectiveness of our approach in identifying the bioactivity of uncharacterized compounds.

## 2 Methods

### General workflow

After preprocessing the microscopy images (optional), our workflow is divided into two main parts, as shown in Figure 2. In the first part, we focus on learning representations of drug-induced perturbations on cells. This is achieved by calculating representations from Cell Painting image data using a semi-supervised contrastive learning approach. Each epoch involves training the CNN in a self-supervised manner on the entire dataset, using the multiple views and replicates of data points to define positives. Concurrently, during the same epoch, the CNN undergoes training with supervised contrastive learning on the annotated subset, where the MeSH classes are used to define positives.^39^ No decision function or classifier is learned during the supervised contrastive learning phase. The annotations are solely used to define the positives.

**Figure 2.**
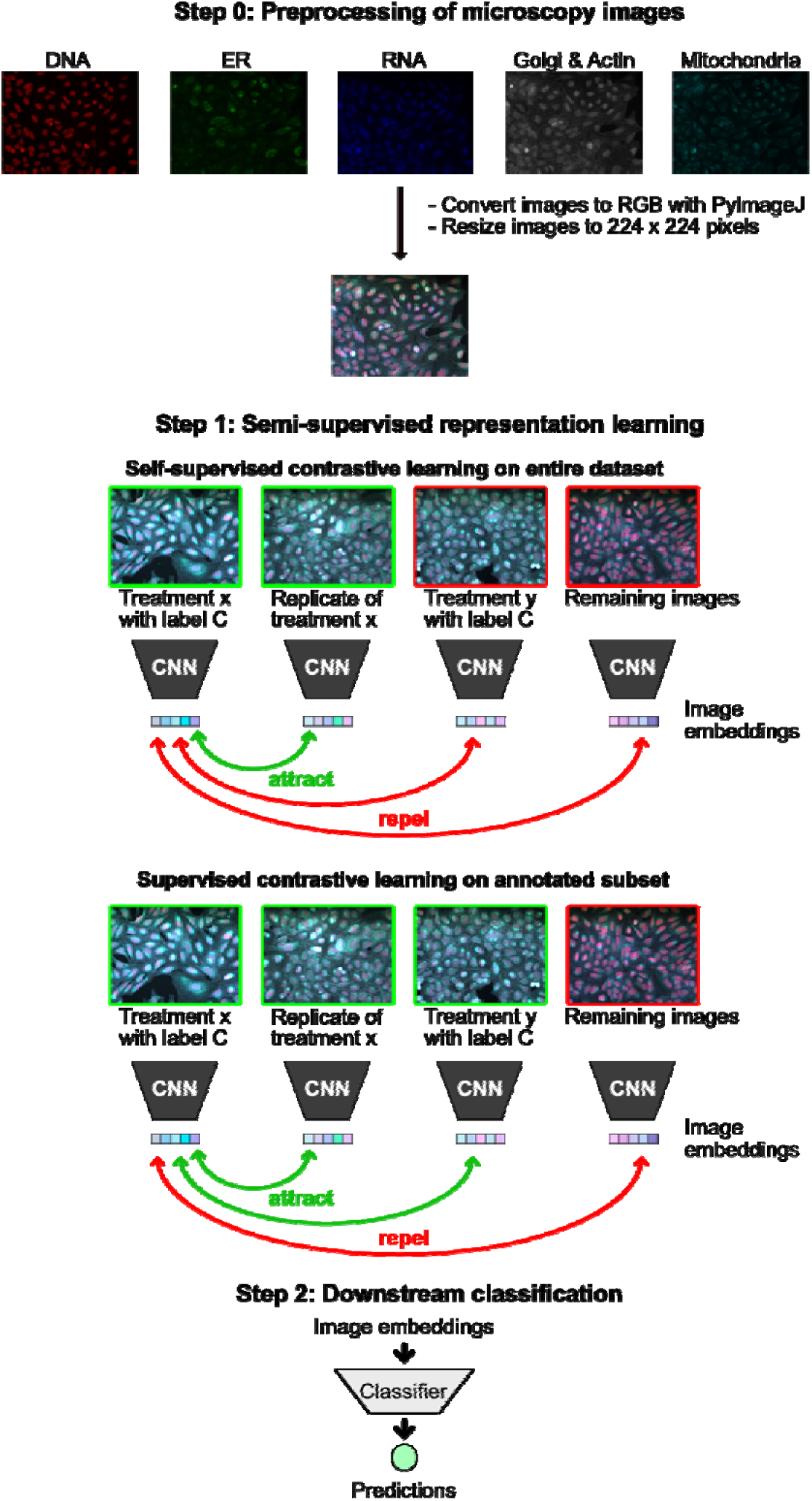
Schematic overview of the SemiSupCon workflow trained on microscopy images. The images were pre-processed by converting the 5-channel microscopy images into RGB images using PyImageJ and they were then resized to 224 x 224 pixels. Then semi-supervised contrastive learning is used to learn image representations of the microscopy images. During contrastive learning, a minibatch of images is sampled, with one image where compound x was used as treatment serving as the anchor. In the self-supervised learning, a well or batch replicate of this image is selected as a positive, while all other remaining images in the batch are designated as negatives. In the subsequent supervised contrastive learning phase, using an annotated subset, all images sharing the same bioactivity label as the anchor are designated as positives. In the last step, these learned representations are used to predict the biological annotations of the compounds using downstream machine learning models.

In the second part, the learned representations are evaluated by using them as input features for downstream machine learning classifiers, aimed at predicting the bioactivity of the compounds. We utilized multiple biological annotation systems for this evaluation. The trained models were then used to predict MeSH classes of unannotated compounds, and the contrastive loss was calculated for each batch. Compounds were then ranked based on the contrastive loss calculated batchwise. Finally, we evaluated the predictions for the compounds with the lowest contrastive loss through literature searches, to confirm the validity of the predicted MeSH classes.

### Dataset

We utilized two publicly available Cell Painting datasets from the Broad institute, BBBC022^40^ and BBBC036^41^, wherein U2OS cells were treated with a large number of bioactive compounds.^42^ The images were captured across five fluorescent channels to analyze various cellular components, including the nucleus, endoplasmic reticulum, mitochondria, nucleoli, F-actin, Golgi apparatus and plasma membrane. All images were taken at 20x magnification, capturing nine sites per well. The BBBC022 dataset includes 69,084 microscopy images (9,216 of which are from the DMSO control), with 1,600 compounds used for treating the U2OS cells. The BBBC036 dataset which we used for training models contains 916,961 images, including 158,859 images from the DMSO control, and it involved the treatment of cells with 30,616 different compounds.

### Annotations

In this study, we used three types of bioactivity annotations: (i) pharmacological classes known as MeSH^37^ classes from PubChem; (ii) MoA and (iii) target annotations, both of which were collected from the Drug Repurposing Hub of the Broad Institute^43^. A total of 839 MeSH classes were extracted from PubChem for the BBBC022 dataset, covering 298 different MeSH classes, with 52 % of all compounds annotated. For the BBBC036 dataset, we obtained 1,079 MeSH classes from PubChem, encompassing 317 MeSH classes, with 4 % of all compounds annotated. It is worth noting that a single compound can be associated with multiple MeSH annotations in PubChem. In the BBBC022 dataset, 498 compounds had multiple classes, whereas in the BBBC036 dataset, 615 compounds fell into this category. Additionally, the MoAs & targets annotations of the compounds were collected from the Drug Repurposing Hub. In the BBBC022 dataset, 773 compounds had MoA annotations with 291 different classes, and 80 compounds had multiple labels. 587 compounds had target annotations with 774 different classes, and 360 compounds had multiple labels. A more detailed description of the annotations can be found in the supplementary (Table S1).

### Setup

Preprocessing: Neural networks were trained using either the 5 fluorescence channel TIFF images directly or RGB images. When training on RGB images, the 5 fluorescence channels of the microscopy images were converted to 3-channel RGB images using PyImageJ^44^. The images were then resized from their original dimensions of 520×696 pixels to a size of 224×224 pixels, for training contrastive learning-based models. Neural networks were trained with the RGB images unless otherwise specified.

### Training

ResNet50^45^ was used as the backbone for the contrastive learning models. The SupContrast^39^ library was used to compute both the contrastive loss and the supervised contrastive loss. The learning rate was set to 10^-2.5^ and the temperature to 0.07. The feature vector size of the projector network was set to 224. Instead of using augmented images, we sampled replicates of the images as positives (see Figure 2). For supervised contrastive learning, MeSH class labels were additionally used to define positives during neural network training. In instances where compounds were associated with multiple MeSH classes, only the first MeSH class was taken into account. In prior research in this field, deep learning models were often trained in a transductive manner^25^, since the main goal is to learn suitable representations for the treatments in a specific cell painting dataset. In our work we trained the deep learning models in a transductive semi-supervised manner, where we incorporate annotations for learning suitable representations. This strategy has shown in several computer vision applications^46–48^ and genome analysis studies^49,50^ to be a data-effective approach, leveraging existing annotations to infer labels of unannotated data points.

The batch size was chosen to be as large as possible, with the GPU RAM of our GPUs determining the maximum batch size. We primarily used 4 NVIDIA A16 GPUs for our calculations (batch size 40). Additionally, calculations were performed with 3 NVIDIA A40 GPU cards (batch size 30). Our contrastive learning-based strategies were trained with a batch size of 40 unless otherwise specified. The models were trained for up to 250 epochs. To assess whether the models converged, both the contrastive loss and downstream MeSH classification were evaluated using a random forest trained with 80 % of the images. The PyTorch^51^ framework was used to implement all the neural networks in this work.

### 2.1 Representation learning

#### Contrastive loss

A lower-level representation of the images was calculated via semisupervised contrastive learning as proposed by Khosla *et al*.^39^ Every minibatch contains N images and for each image x a replicate x∼ is randomly selected as sample pair, so that there are 2N images in total inside the batch. The set of 2N samples will be from now on referred to as “multiviewed batch”, whereas the set of N sample pairs will be referred to as “batch” from now on. ResNet50 was used as encoder network denoted by f and a multi-layer perceptron (MLP) was used as the projector head denoted by g to get the latent representation z= g(f(x)). In this work we used z as the representation of the images.

An image inside of the batch is chosen as anchor, to which the remaining images are contrasted against. Let iE I = {1… 2N} be the anchor index of an arbitrary sample within a multiviewed batch, let j(i) be the index of the corresponding replicate (positive) and the remaining 2(N-1) indexes A_(i)_ belong to the negatives. The self-supervised loss L^self^ is then defined as follows:

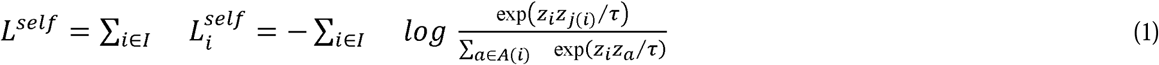

where *z_i_* and *z_j_* are the latent representations of the positive pair, *z_a_* is the latent representation of a negative sample and τ is a scalar temperature parameter. To extend the contrastive loss to the supervised contrastive loss, where multiple samples can be designated as positives if they belong to the same class as the anchor, the following supervised contrastive loss equation L^sup^ from Khosla *et al*.^39^ was used:

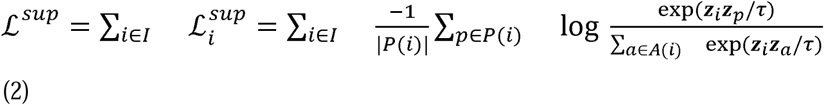

The indices of the positives of the anchor i, which belong to the same class are denoted as P(i) and |P(i)| is the number of positives in the multi viewed batch. This supervised contrastive loss allows us to incorporate knowledge about biological annotations, to learn suitable image representations. Since the SupContrast library only supports one label for each compound we only used the very first annotation of the compounds for supervised contrastive learning, the remaining annotations were not considered. This is a limitation of this supervised contrastive learning approach, which will be addressed in future works. Only the MeSH classes were used in this work as labels during supervised contrastive learning. After learning the image representations, downstream classification was performed to evaluate if the learned representations were able to capture the pharmacological similarity between the different compounds in the dataset.

### 2.2 Downstream analysis

Downstream bioactivity prediction was performed with image-level features which were calculated with semi-supervised contrastive models. These results were compared with multiple methods for calculating phenotypical representations of cell painting image data, which have been previously evaluated in the literature for bioactivity prediction. Baseline methods for calculating the representations included a self-supervised contrastive model (comparable to Perakis *et al.*^25^), DINO^29^ and profiles derived from CellProfiler. The representations were used as features in downstream random forest (RF) and multi-layer perceptron models to predict the biological annotations of the data points.

BBBC022 served as the baseline dataset for downstream bioactivity prediction. Additionally, we evaluated bioactivity prediction on the BBBC036 dataset. However, due to the relatively large size of the dataset we were only able to perform a limited number of calculations on this dataset. For BBBC036, we only calculated representations using the semi-supervised contrastive learning strategy and DINO using RGB images, as training was faster compared to using the 5-channel TIFF images.

For DINO, we used the same architecture and setup as Doron *et al.*^29^, except that we used weaker augmentations. Similar to SemiSupCon, we evaluated models trained with both TIFF images and RGB images on the BBBC022 dataset. For the model trained with RGB images, we used the same augmentations as Doron *et al*. except for random brightness and contrast changes. For the model trained with TIFF images, we used only random cropping, vertical and horizontal flipping as augmentations. We evaluated a MLP with 3 layers as downstream classification model, where the first layer had 512 hidden units and the second had 256 hidden units, as described by Doron *et al*.^29^ Rectified linear units (ReLU) served as the activation function for the input and hidden layers, and dropout was applied to the first two layers (with a probability set to 50 %).The learning rate was set to 10^-2.5^ and cross entropy was chosen as the loss function. Early stopping was used during the training of the downstream MLPs.

Downstream bioactivity prediction was evaluated using both single-label and multi-label classification tasks. For multi-label classification, multi-layer perceptrons and random forests were used to evaluate prediction of MeSH classes, Drug Repurposing Hub MoA and target annotations. Single-label MeSH class prediction was evaluated only with random forest classifiers. In our evaluations, 80% of the treatments from the annotated subsets were allocated for training, while the remaining 20% were reserved for evaluation and calculation of metrics. Throughout our evaluation of various strategies for calculating image representations, the downstream analysis remained consistent. To validate our downstream models, 5-fold cross validation (5-CV) was used. This involved creating 5 different training and test splits of our dataset, ensuring that each data point was used at least once for training and testing. A fully supervised model was also evaluated as baseline for multi-label MeSH class prediction. ResNet50 was used as the backbone for the supervised model and the architecture of the classification head was designed to ensure comparability with the contrastive learning-based approach. The classification head corresponded to a combination of the projection head and downstream MLP, similar to what was used in the contrastive learning-based strategies.

#### Evaluation metrics

For classification, accuracy and the averaged area under the precision-recall curve (PR AUC) were used as evaluation metrics. The PR AUC is calculated for each individual class and then averaged across classes. The accuracy for the single-label and multi-label predictions is calculated as the mean over all predictions on the test set that matched the provided annotations.^52^

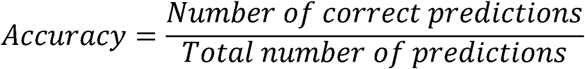

#### Number of correct predictions Total number of predictions

For the calculation of the multi-label accuracy, predictions were only considered as correct if all predicted labels of a data point match the ground truth annotations.

We calculated statistical reports to evaluate the performance of our workflow for each individual MeSH class. Our objective was to analyze the MeSH classes for which our workflow performs particularly well and those where it does not exhibit any predictive ability. Here, we employ precision, recall, and the F1-score as evaluation metrics.

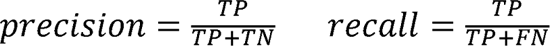

TP are the true positives, TN true negatives, FN false negatives and FP are the false positives. The F1-score represents the harmonic mean between precision and recall in a classification problem:

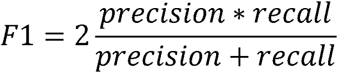

The potential of heat-diffusion for affinity-based trajectory embedding (PHATE)^53^ algorithm was used to visualize the phenotypic space.

## 3 Results

### 3.1 Evaluation on downstream tasks

We evaluated the image representations generated by our semi-supervised contrastive learning strategy (SemiSupCon), against those generated from a self-supervised contrastive learning strategy (Con), DINO, and expert-engineered profiles by CellProfiler, which served as baseline comparisons for downstream bioactivity prediction. This evaluation aimed to determine whether the corresponding representations are able to capture the pharmacological similarity between different compounds by assessing their ability to correctly predict the bioactivity labels of annotated compounds. We used MeSH classes from PubChem, as well as MoA and biological target annotations from the Drug Repurposing Hub, as labels for multi-label classification of compounds from the BBBC022 dataset. For this baseline comparison we used a downstream MLP with the same architecture as described by Doron *et al*.^29^

Utilizing the features derived from SemiSupCon increased downstream bioactivity prediction accuracy and PR AUC across all downstream tasks when compared to the baselines (Figure 3). SemiSupCon attained an accuracy of 21.0 % for the prediction of MeSH classes, while the other strategies yielded multi-label accuracies of zero or close to zero. The highest predictive performance among the baselines was achieved with the Con strategy. DINO and CellProfiler exhibited zero accuracy in predicting any of the Drug Repurposing Hub annotations, whereas Con achieved accuracies of 3.4 % for Drug Repurposing Hub MoA classification and 1.0 % for target classification. SemiSupCon improved downstream prediction accuracies compared to the baselines by 8.4% for Drug Repurposing Hub MoA annotations and 2.5% for target annotations. Although the SemiSupCon strategy yielded considerably lower accuracies and averaged PR AUC values when classifying Drug Repurposing Hub annotations compared to MeSH classes, it still demonstrated higher predictive performance compared to the baselines. This suggests that the learned representations of SemiSupCon can generalize to new classification tasks. To further evaluate the performance of the representations learned by SemiSupCon, we trained an additional configuration, this time adopting an inductive strategy for SemiSupCon. We also included a fully supervised ResNet50 model as baseline, which was evaluated for multi-label MeSH class prediction. The results showed zero accuracy for the fully supervised model (Table S2), indicating that the annotated subset is too small to properly train a supervised model. The inductive SemiSupCon strategy slightly improved downstream prediction of MeSH classes and Drug Repurposing Hub MoA annotations compared to the baselines (Table S2).

**Figure 3.**
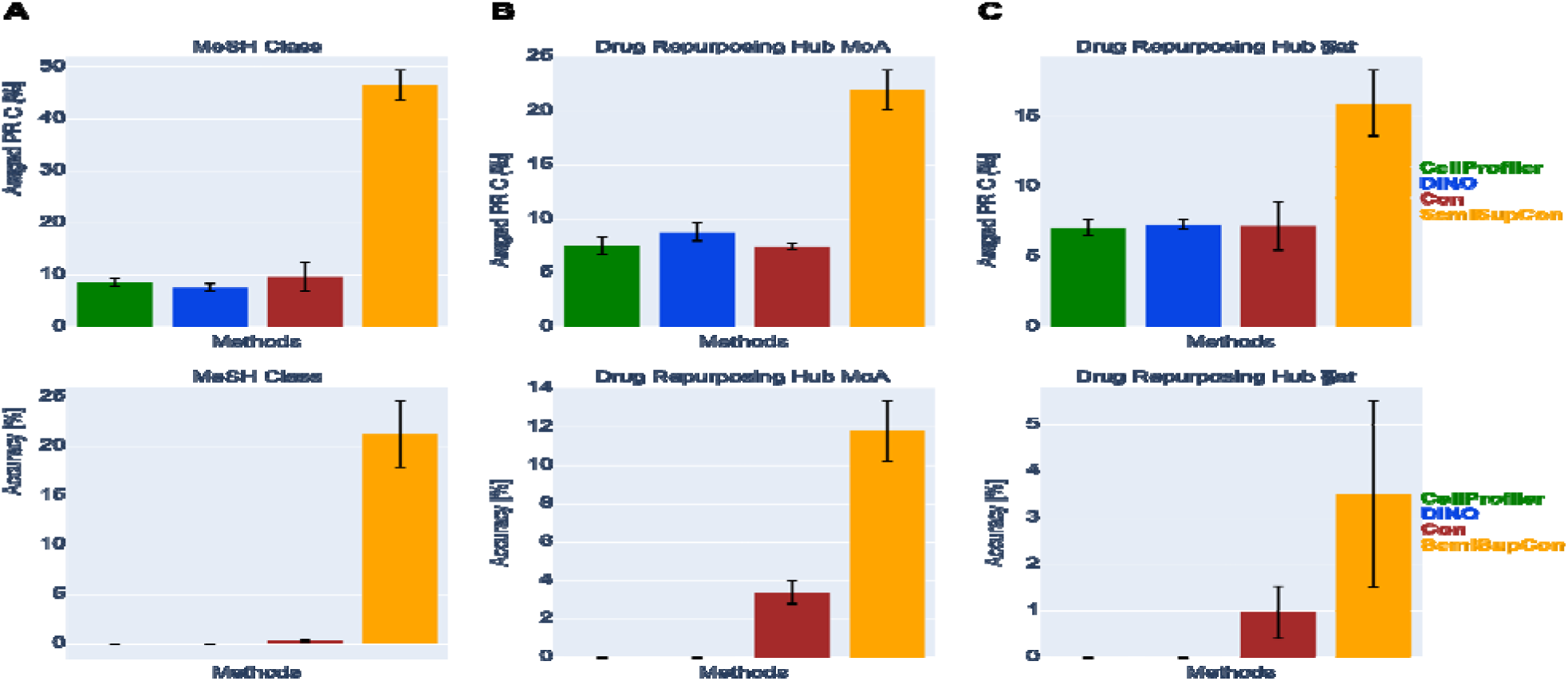
Baseline comparison for downstream bioactivity prediction on the BBBC022 dataset. Deep learning-based features and CellProfiler calculated features were evaluated for predicting (A) MeSH class annotations and annotations from the Drug Repurposing Hub as labels, (B) MoAs and (C) biological targets. Features calculated with CellProfiler (green), DINO (trained on TIFF images; blue), Con (brown), and SemiSupCon (orange) were used as inputs for training MLPs for multi-label classification. Averaged PR AUC (top) and accuracy (bottom) were used as metrics. 5-CV was used for this evaluation and the results were averaged across the different folds.

Additionally, we evaluated several strategies and configurations for downstream classification on the BBBC022 and BBC036 dataset using RFs as classifiers. Image representations calculated with CellProfiler, DINO, Con and the SemiSupCon strategy were evaluated. RFs were trained on multi-label tasks to predict the MeSH classes as well as Drug Repurposing Hub MoA and target annotations. Additionally, a RF model was trained on a single-label task to predict the first MeSH class of the compounds. Accuracy was used as metric for these evaluations. The results for the downstream classification on the BBBC022 dataset are summarized in Table 1.

**Table 1.**
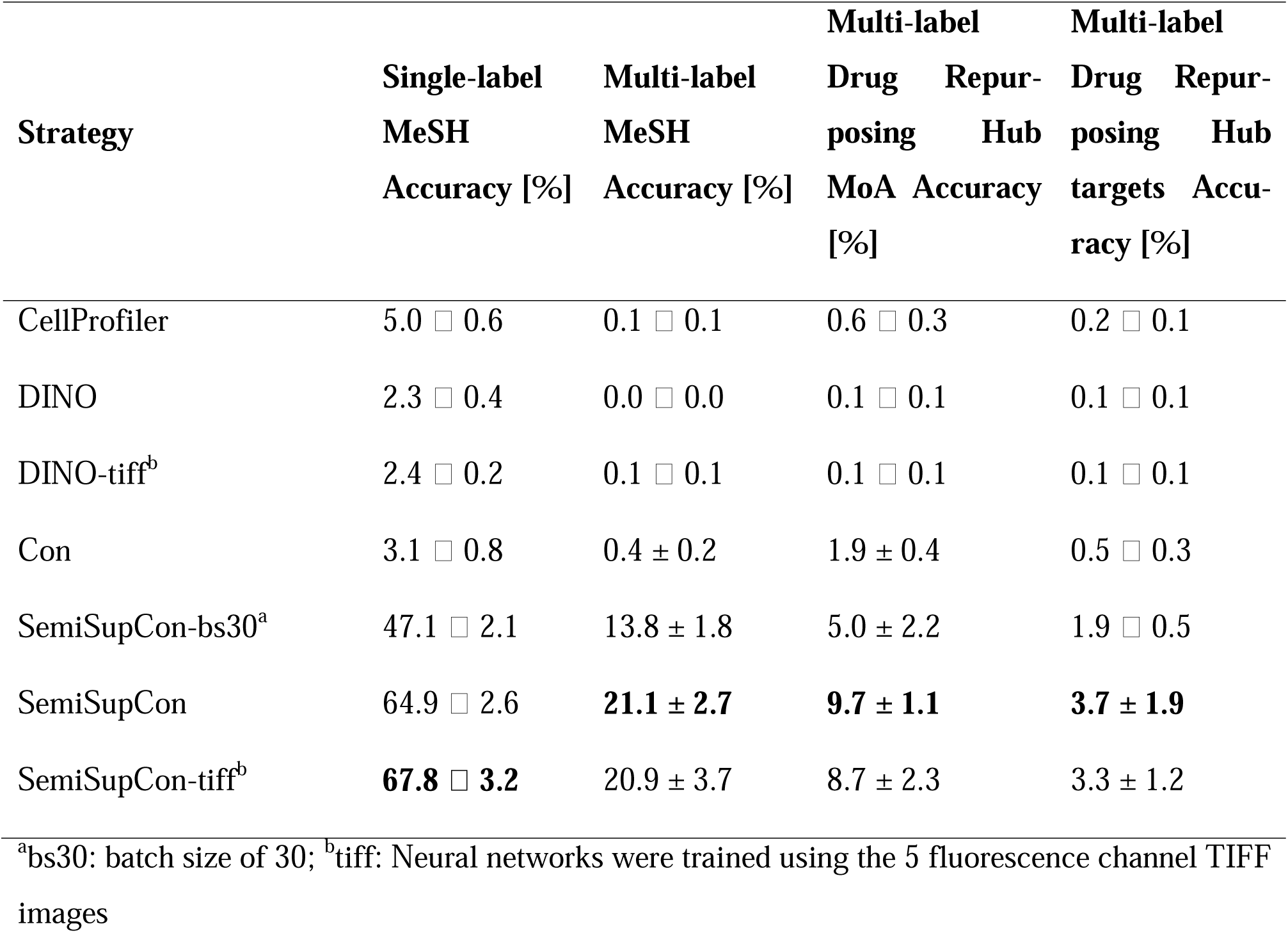
Comparison of downstream classification with RFs across five cross-validation folds on the BBBC022.

SemiSupCon also exhibited improved downstream classification using RFs compared to the other methods (Table 1). Using a smaller batch size with SemiSupCon-bs30 decreased downstream performance on all tasks. The Con strategy attained higher mean accuracies across all multi-label downstream tasks compared to the other baselines.

The results for downstream classification with RFs using representations from SemiSupCon and DINO on the BBBC036 dataset are summarized in Table 2. Notably, the SemiSupCon strategy achieved a much lower accuracy on the BBBC036 dataset, compared to its performance on the BBBC022 dataset, despite the BBBC036 dataset containing more than 10 times the number of images. When applied to the BBBC036 dataset, the SemiSupCon strategy demonstrated considerably higher accuracy in single-label MeSH class prediction compared to DINO, although its performance in multi-label prediction was only slightly better overall.

**Table 2.**
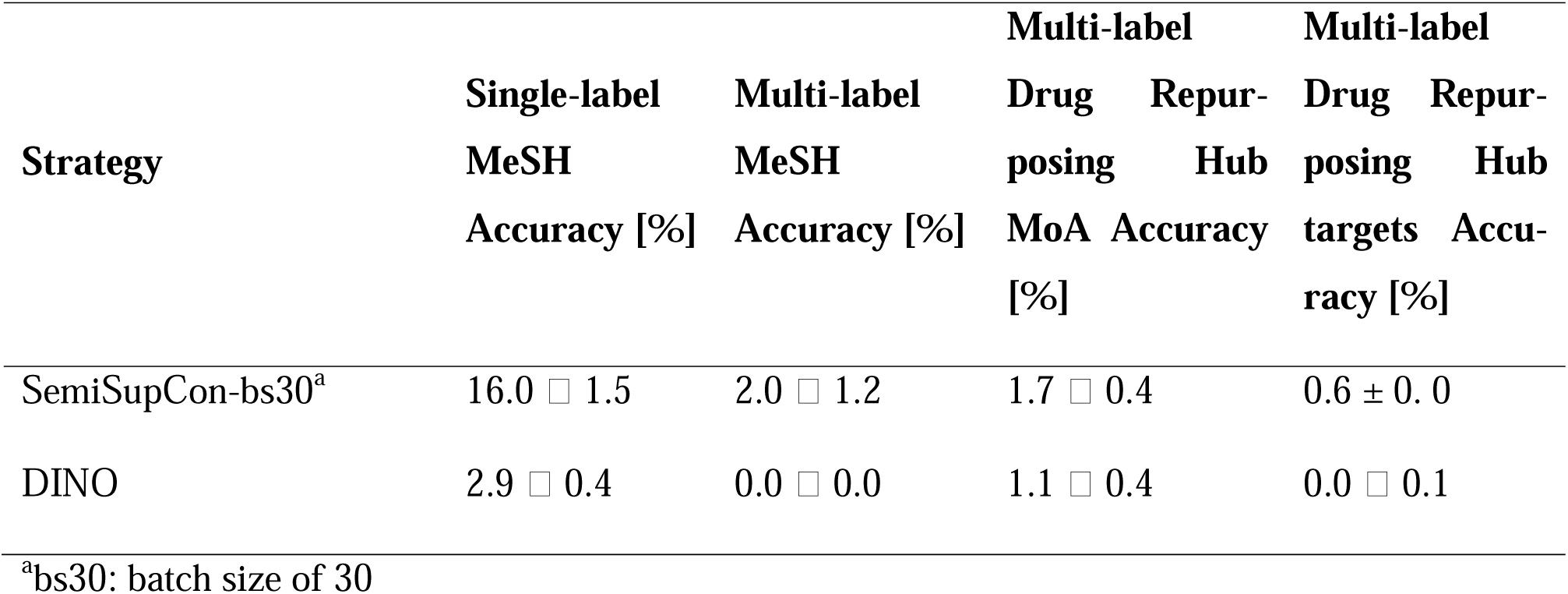
Comparison of downstream classification with RFs across five cross-validation folds on the BBBC036 dataset.

In conclusion, comparing bioactivity prediction between SemiSupCon and the baselines demonstrates the utility of using SemiSupCon-based image features to represent compound perturbations in the Cell Painting assay and for bioactivity prediction.

### 3.2 Phenotypical space exploration

As the next step, we visualized the learned phenotypical space of the SemiSupCon(BBBC022) model using the PHATE algorithm (Figure 4; for comparison, visualization using the uniform manifold approximation and projection (UMAP) algorithm is shown in Figure S1), as this model achieved the highest single- and multi-label accuracy among different methods (Table 1). In this way compounds with similar bioactivities can be identified without the need for biological annotations. The results, as shown in Figure 4A, reveal that the DMSO embeddings are clearly separated from the other data points in the phenotypical space. This distinct separation is anticipated since DMSO serves as the negative control and is overrepresented in the dataset. Additionally, we focused on visualizing the phenotypical space of the MeSH class-annotated subset (Figure 4B). In this visualization, we can observe multiple regions where treatments sharing the same MeSH class cluster together in the phenotypical space (Figure 4C and Figure 4D for close-up view of two such regions), suggesting that the semi-supervised approach effectively learns to associate treatments with similar bioactivities. For comparison, PHATE visualization of the CellProfiler profiles calculated from the BBBC022 dataset is shown in Figure S2.

**Figure 4.**
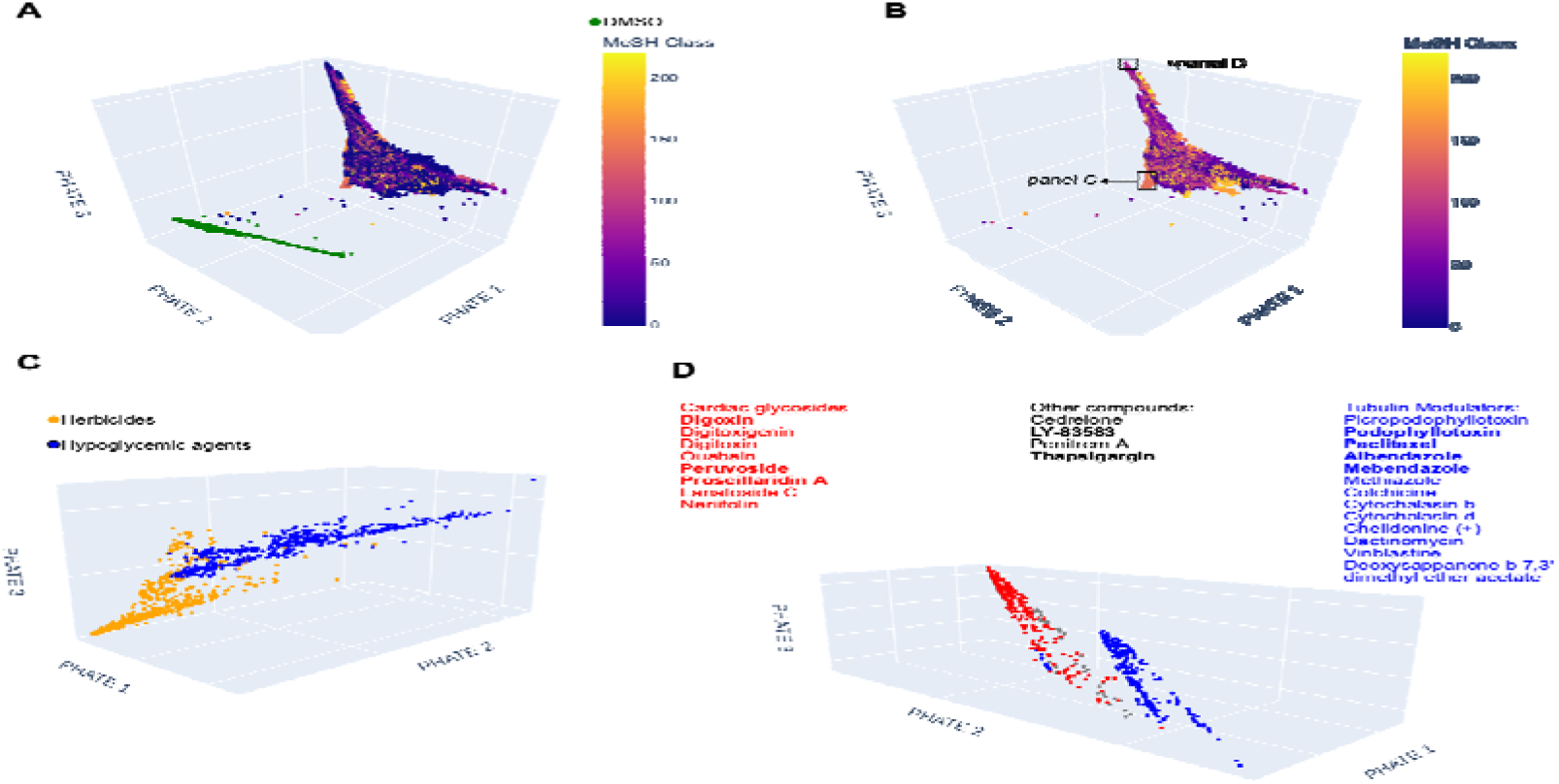
3D visualization of the phenotypical space from SemiSupCon(BBBC022) features, calculated with the PHATE algorithm. The phenotypical space was explored to identify regions where compounds with similar bioactivities form clusters. (A) Visualization of all data points, colored according to their MeSH classes. Compounds with multiple MeSH classes, were assigned to their first MeSH class. Data points with the lowest color code (0) indicate the unavailability of a MeSH class, while green data points represent the DMSO controls. (B) Visualization focusing on the subset of data points annotated with MeSH classes. (C) Close-up view of the phenotypical space, showing only herbicides and hypoglycemic agents. Literature searches of compounds in this region revealed plausible reasons for the proximity of these two classes in the phenotypical space, as several compounds were found to belong to both. (D) Close-up view of all compounds at the upper peaks of the phenotypical space revealed cardiac glycosides and tubulin modulators clusters. Compounds confirmed as cardiac glycosides or tubulin modulators through literature searches are listed, with those containing corresponding MeSH annotations highlighted in bold.

Analyzing the phenotypical space, we found that herbicides form a clear and distinct cluster in this space (Figure 4B and 4C), showing very close proximity to several hypoglycemic agents. A literature search of compounds at the interface between these two clusters reveals a biologically plausible explanation for the proximity of these two pharmacological classes. Several hypoglycemic agents are organic compounds belonging to the family of sulfonylureas (e.g. Chlorpropamide), known in the literature for their usage as herbicides.^54^ 2,4,5-trichlorophenoxyacetic acid is a non-sulfonylurea herbicide, and although no direct evidence was found to classify this compound as a hypoglycemic agent, but its close derivative, 2,4-dichlorophenoxyacetic acid, serves both as herbicide and a hypoglycemic agent.^55^

We also identified clusters for tubulin modulators and cardiac glycosides cluster at the peaks of the learned phenotypical space (Figure 4D), without relying on the MeSH classes. The two clusters were well separated yet in close proximity to each other and a large proportion of the compounds in those regions of the phenotypical space, were confirmed to belong to the corresponding class of compounds. The cardiac glycosides cluster comprised structurally related compounds, including the following cardenolides: digoxin^56^, digitoxin^57^, digitoxigenin (an aglycon of digitoxin)^58^, ouabain^59^, peruvoside^60^, proscillaridin A^61^, lanatoside C^62^ and neriifolin^63^. Only a few of these compounds had MeSH class annotations, highlighting the capability of the learned representations to capture the pharmacological similarity of unannotated compounds. It should be noted that there is no “Cardiac Glycosides” MeSH class; however, annotated cardiac glycosides commonly have the MeSH class “Cardiotonic Agents”.

The tubulin modulator cluster contained many structurally related compounds. A literature search confirmed the following compounds as modulators of tubulin: picropodophyllotoxin^64^, podophyllotoxin^64^, paclitaxel^65^, albendazole^66^, mebendazole^67^, colchicine^68^, cytochalasin b^69^, cytochalasin d^69^, chelidonine (+)^70^, dactinomycin^71^, vinblastine^72^ and deoxysappanone b 7,3’-dimethyl ether acetate^73^. Most MeSH class annotated compounds in the tubulin modulator cluster had the MeSH class “Tubulin Modulator”. Four compounds in Figure 4D could not be identified as either cardiac glycosides or tubulin modulators. The MeSH class annotated compounds among them commonly had the MeSH class “Enzyme Inhibitors”. It is worth noting that the cardiac glycosides and tubulin modulator clusters were also identified in the publication of the original BBBC022 dataset by Gustafsdottir *et*. *al*.^42^, where CellProfiler features were clustered using hierarchical clustering.

### 3.3 Analysis of MoA classifications

Predictions from downstream RF trained with SemiSupCon(BBBC036) and SemiSupCon(BBBC022) representations were further analyzed, as these two strategies achieved the highest accuracy in their respective datasets. To this end, we performed a deconvolution of the predictions by statistically evaluating the predictive performance for each MeSH class. In our previous evaluation of downstream classification, the predictive performance was averaged across all classes, the goal of this analysis however was to discern which individual MeSH classes can be effectively predicted by our approach and which not. A 5-CV was conducted to allow all data points to be analyzed in this evaluation by aggregating all the test sets from the 5 folds together. We calculated the mean precision, recall and F1-score across all folds.

Out of 223 MeSH classes, 106 were predicted with a precision higher than 10 %, using a RF trained with single-label MeSH classes on SemiSupCon(BBBC022) features. The top 25 MeSH classes with the highest F1-scores are summarized in Table 3. Notably, for most of these top 25 MeSH classes, a high F1-score was achieved, with the majority displaying an F1-score above 0.90. The mean precision over all 223 MeSH classes was 81.48 % and the mean recall was 43.25%.

**Table 3.** Predictive performance for 25 MeSH classes with highest F1-score based on the single-label RF trained on SemiSupCon(BBBC022) features.

| MeSH class | Precision [%] | Recall [%] | F1-score |
| --- | --- | --- | --- |
| Hypoglycemic Agents | 99.33 | 99.45 | 0.99 |
| Antihypertensive Agents | 99.72 | 98.05 | 0.99 |
| Antipsychotic Agents | 99.46 | 97.80 | 0.99 |
| Anti-Infective Agents | 99.54 | 97.65 | 0.99 |
| Anti-Bacterial Agents | 99.83 | 98.36 | 0.99 |
| Dopamine Agonists | 99.35 | 96.16 | 0.98 |
| Antifungal Agents | 99.31 | 96.03 | 0.98 |
| Serotonin Antagonists | 99.44 | 96.05 | 0.98 |
| Calcium Channel Blockers | 99.26 | 95.84 | 0.98 |
| Anti-Inflammatory Agents | 98.18 | 97.45 | 0.98 |
| Anti-Inflammatory Agents, Non-Steroidal | 99.13 | 96.95 | 0.98 |
| Vasodilator Agents | 99.08 | 97.82 | 0.98 |
| Herbicides | 99.21 | 95.18 | 0.97 |

| **MeSH class** | **Precision [%]** | **Recall [%]** | **F1-score** |
| --- | --- | --- | --- |
| Diuretics | 99.35 | 94.08 | 0.97 |
| Enzyme Inhibitors | 98.80 | 94.59 | 0.97 |
| Dopamine Antagonists | 98.66 | 94.09 | 0.96 |
| Anti-Arrhythmia Agents | 99.35 | 92.39 | 0.96 |
| Anti-Allergic Agents | 96.25 | 93.64 | 0.95 |
| Insecticides | 98.43 | 90.76 | 0.94 |
| Antiemetics | 96.67 | 91.27 | 0.94 |
| Cholinesterase Inhibitors | 96.98 | 87.68 | 0.92 |
| Adrenergic alpha-Agonists | 99.17 | 84.72 | 0.91 |
| Anesthetics, Local | 99.44 | 78.68 | 0.88 |
| Bronchodilator Agents | 99.16 | 77.84 | 0.87 |
| Anthelmintics | 98.61 | 75.51 | 0.86 |

When analyzing the results from Table 3 we observe that the predictive performance for the top 25 MeSH classes is much higher than the other classes. We conducted a similar analysis using the single-label RF model trained on SemiSupCon(BBBC036) features (Table 4). Here, the predictive performance for individual MeSH classes is significantly lower compared to the results from the BBBC022 dataset. This reduction in the predictive ability of individual MeSH classes is anticipated, given that the overall accuracy of the RF on the BBBC036 dataset is much lower, as shown in Table 2. Nonetheless, there is a notable overlap of 64% for the top 25 MeSH classes when comparing these two datasets. Given that the models were individually trained on two different CP datasets, these findings strongly suggest that a substantial number of pharmacological MeSH classes can be identified with sufficient precision using the proposed workflow. The mean precision over all MeSH classes was 6.05 % and the mean recall was 5.35 %.

**Table 4.**
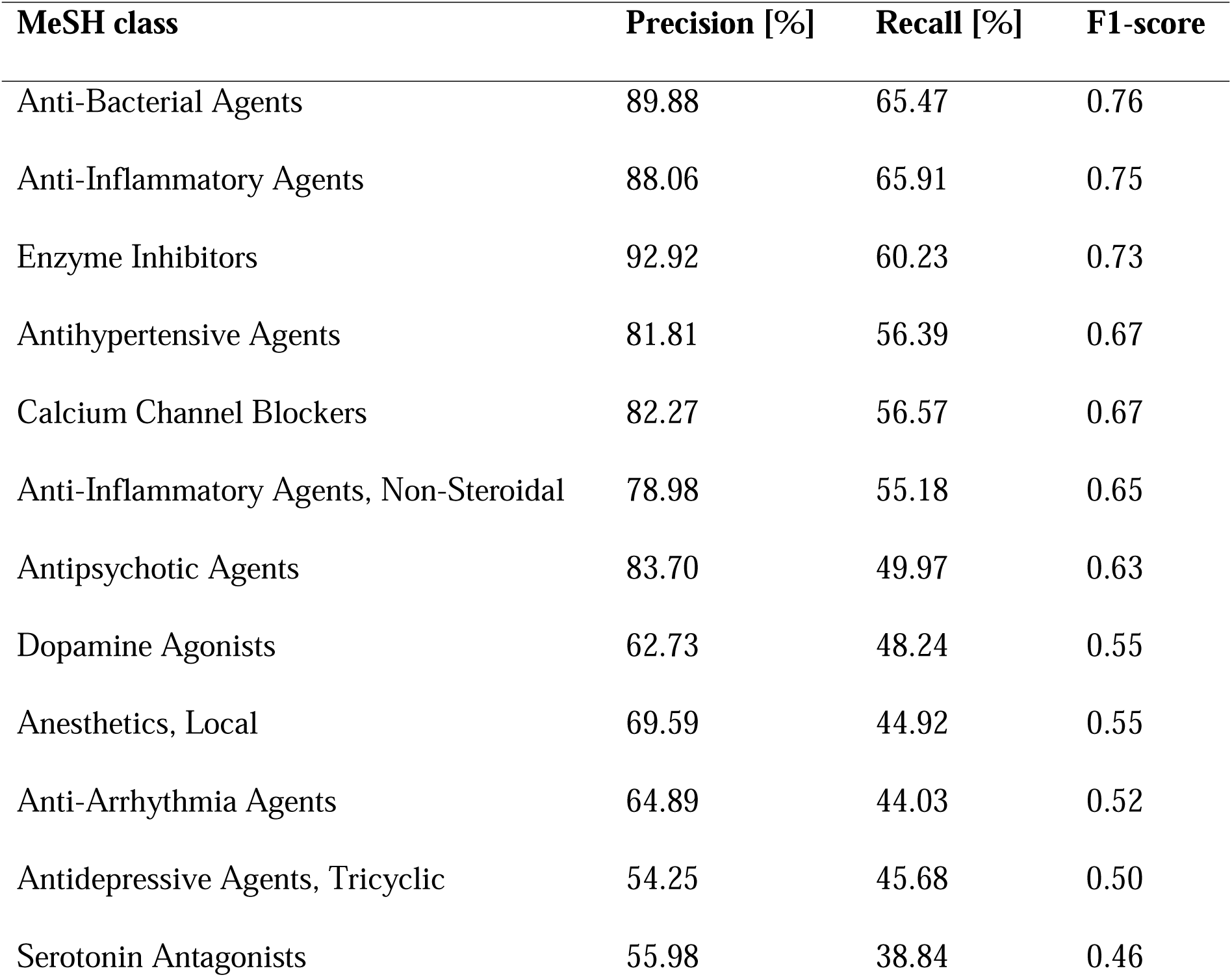
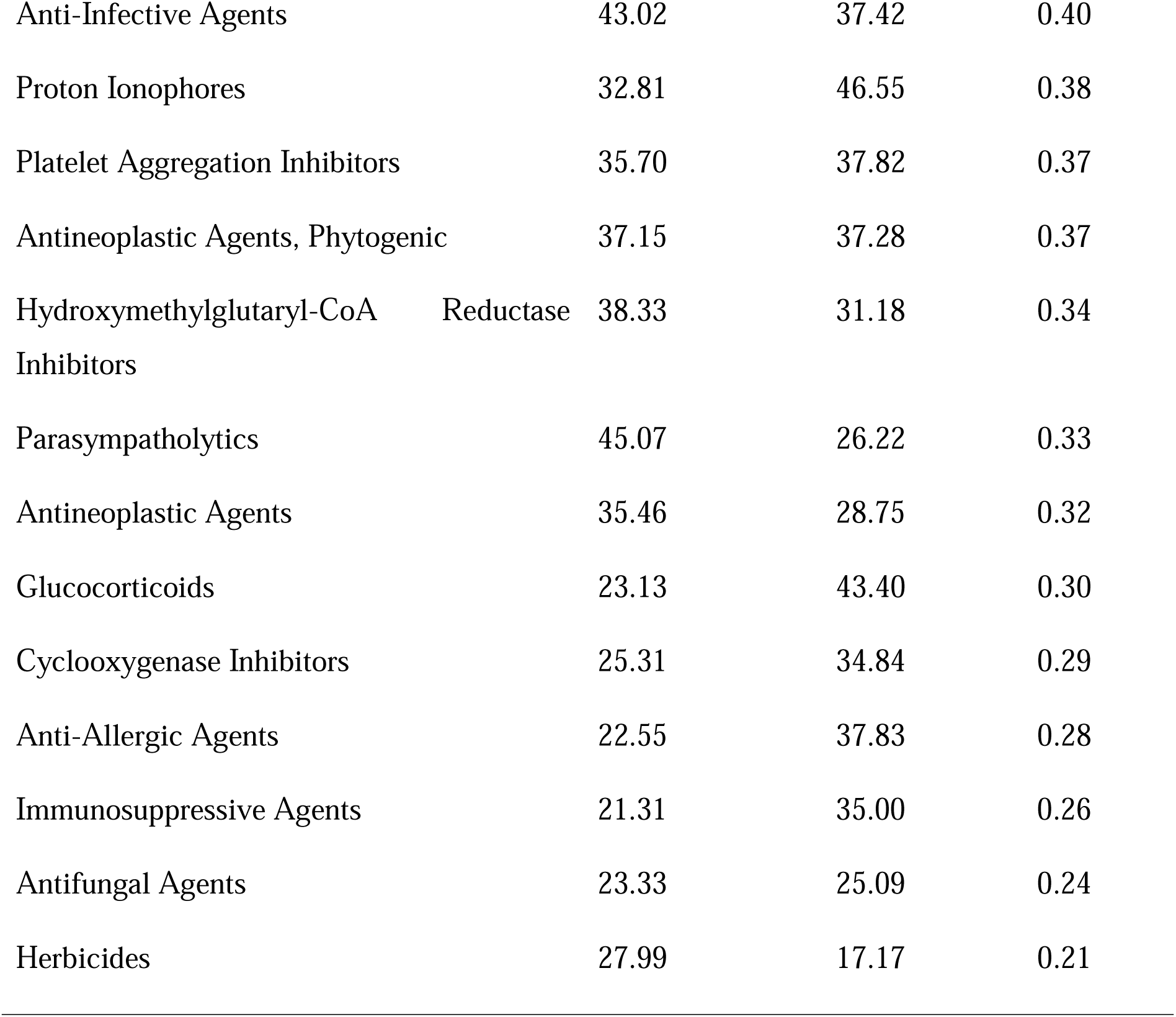
Predictive performance for 25 MeSH classes with highest F1-score based on the single-label RF trained on SemiSupCon(BBBC036) features.

Downstream classification on the BBBC036 dataset showed much lower predictive performance, compared to the BBBC022 dataset, which is also reflected when analyzing the top 25 classes (Table 4). We observed that only the top 3 classes achieve a F1-score higher than 0.7 (in Table 3 lowest F1-score is 0.86). The F1-score is than rapidly decreasing for the remaining classes, with the lowest F1-score in the Table 4 being 0.21.

### 3.4 MoA classification of unannotated compounds

Next, we aimed to predict the MeSH classes of unannotated compounds in the BBBC022 dataset using a single-label RF trained on SemiSupCon(BBBC022) features. The self-supervised loss of treatments calculated with our SemiSupCon(BBBC022) model was used to assess how consistent the learned representations are in comparison to other treatments. The compound treatments were ranked based on the batch-wise calculated contrastive loss. We selected the 10 compounds with the lowest self-supervised contrastive loss. For each of these compounds, we used the single-label RF to predict the MeSH classes of all the replicates. The predicted MeSH classes of a compound were subsequently ordered by the count of replicates that shared the same predicted MeSH class.

A manual literature search was performed for the predicted MeSH classes. Predictions were only considered as confirmed if evidence supporting the MeSH class prediction was found in the literature and if that class was among the top 3 ranked predictions. The predicted MeSH classes of compounds with confirmed predictions are shown in Figure 5, where up to 3 MeSH classes with the most predictions are shown. The literature search confirmed the bioactivity predictions for 6 of the 10 compounds directly (indicated in blue in Figure 5): paraxanthine as a bronchodilator agent^74,75^ and cardiotonic agent^76^; 6-formylondolo [3,2-B] carbazole as a bronchodilator^77^ and antihypertensive agent^78^; adrenic acid as an antineoplastic agent^79^; betahistine as an anti-inflammatory agent^80^ and vasodilator agent^81^, beta-dihydrorotenone as an antioxidant^82^; hydrocotarnine as a dopamine agonist^83^. Although no direct evidence supported the predictions for naproxol, but naproxen, a structurally closely related compound, was found to exhibit the predicted activities for naproxol, namely as a dopamine antagonist^84^ and serotonin antagonist^84^ (marked in orange in Figure 5). The remaining compounds for which a literature search could not confirm the prediction are listed in Figure S3. For comparison, four predictions of MoAs, confirmed through literature search, for the 10 unannotated compounds with the lowest contrastive loss in the BBBC022 dataset using a multi-label RF model trained on SemiSupCon(BBBC022) representations are shown in Figure S4.

**Figure 5.**
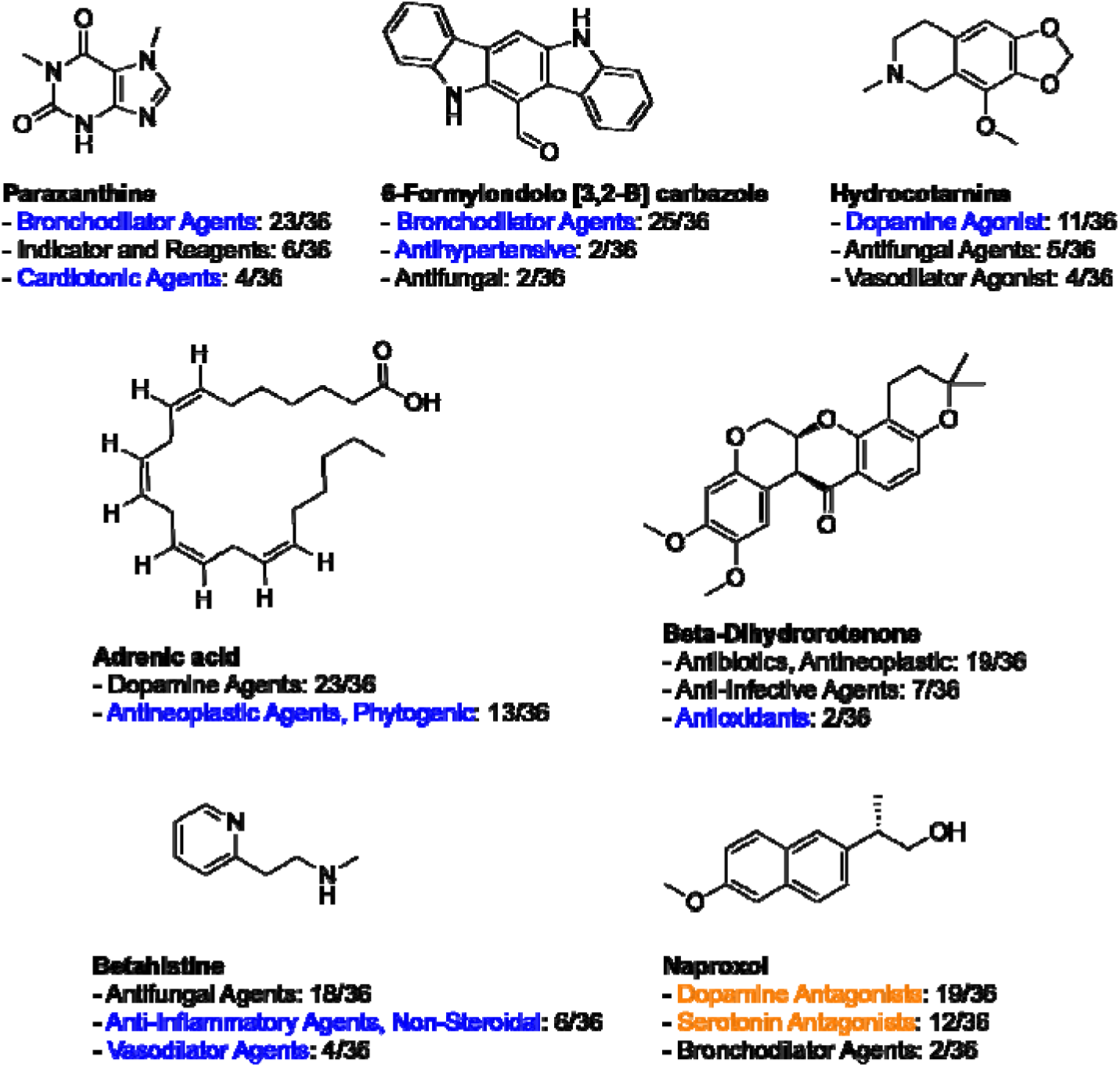
Outcomes of the literature search for 10 unannotated compounds with lowest batchwise-contrastive loss from the BBBC022 dataset. Single-label RF trained on SemiSupCon(BBBC022) features was used to predict the MeSH classes for these compounds. The predicted MeSH classes could be confirmed for 7 molecules through the literature search. Predictions were made for each of the 36 image replicates of a compound, and for each compound, the three most frequently predicted MeSH classes are shown. Predictions that were directly confirmed are indicated in blue, those with indirect evidence from closely related derivatives in orange, and unconfirmed predictions are shown in black.

The same analysis was repeated on the unannotated subset of the BBBC036 dataset, based on predictions from a single-label RF trained on SemiSupCon(BBBC036) representations. Given the generally low downstream accuracy of SemiSupCon(BBBC036) model, we extended our search to include the top 25 unannotated compounds with the lowest batchwise-calculated contrastive loss. Only the replicate of a compound with the lowest contrastive loss was evaluated for this analysis. In this case, we were able to confirm the bioactivity indicated by the MeSH classes for 6 compounds through manual literature searches (Figure 6): BRD-A41941932 as an antihypertensive agent^85^, BRD-K72726508 as an anti-inflammatory agent^86^, BRD-K00732328 as an antipsychotic agent^87^, BRD-A04661934 as an adrenergic alpha-antagonist^88,89^, BRD-A83387524 as a hypolipidemic agent^90^, and BRD-K63608008 as an anti-bacterial agent^91^. For comparison, Figure S5 shows seven MoA predictions that were confirmed through literature searches for the top 25 profiles of unannotated compounds in the BBBC036 dataset with the lowest contrastive loss using a multi-label RF model trained on SemiSupCon(BBBC036) features.

**Figure 6.**
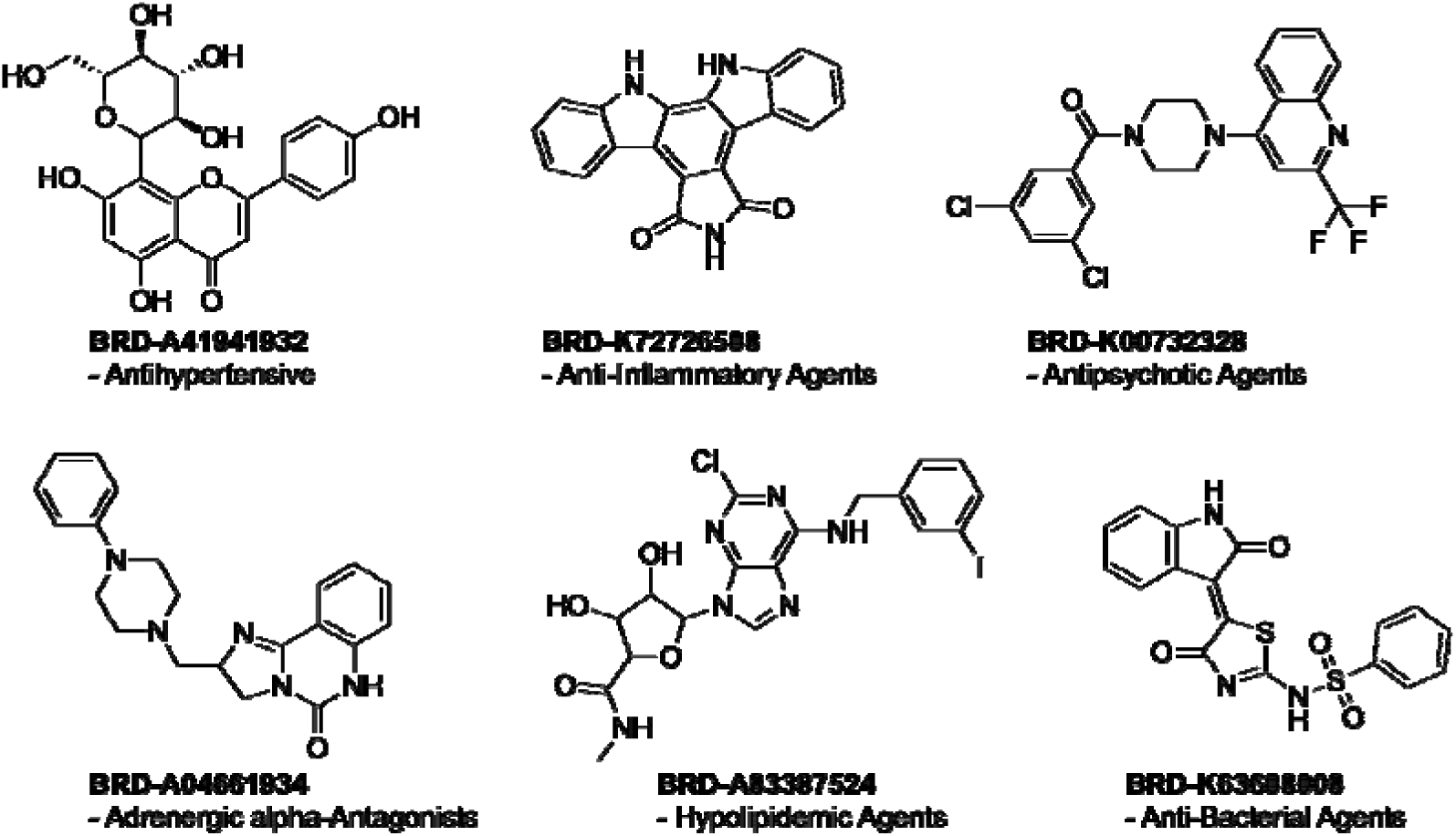
Manual literature searches confirmed 6 out of top 25 unannotated compounds with the lowest batchwise-contrastive loss from the BBBC036 dataset. Single-label RF trained on SemiSupCon(BBBC036) representations was used to predict the MeSH classes.

## 4 Discussion

One important application of the Cell Painting assay, and the profiles generated from it, is to identify compounds with similar phenotypic effects, where similarity to annotated compounds can be used to infer the bioactivity of unannotated compounds.^1^ In this study, we introduced a semi-supervised contrastive learning approach for the bioactivity prediction of small molecules from Cell Painting image data. We leveraged the large amount of image data provided by high-throughput imaging and existing biological knowledge of compounds to learn discriminative representations of the drug-induced perturbations in CP assays. The first MeSH class annotations of the compounds were used to define positives during the supervised contrastive training of SemiSupCon. The phenotypical representations obtained were then used to predict annotations of compounds in various downstream tasks. Our findings demonstrate that SemiSupCon improved downstream classification across three different annotation systems — MeSH classes, Drug Repurposing Hub MoA and target annotations — when compared to expert-engineered representations using CellProfiler and representations based on self-supervised learning methods (Figure 3). This indicates that SemiSupCon is more effective in assessing the similarity of drug-induced perturbations within the phenotypical space, compared to the evaluated baselines.

It should be noted that we evaluated multi-label accuracy, alongside single-label accuracy, which is an overly strict metric as it requires that the predictions for all classes of a query data point match the ground truth annotations. Multi-label accuracy nonlinearly scales the predictive performance of models which can cause sharp and disproportionately large changes when comparing low-performing and high-performing models, making low-performing models appear entirely incapable of performing the multi-label classification tasks.^92^ To more accurately assess the predictive performance of the representations, more labeled data would be necessary, which is a significant challenge due to the scarcity of biological annotations. When a model achieves a much higher average PR AUC compared to the multi-label accuracy, it may indicate that many false predictions are being made for certain classes. For most multi-label classification tasks, SemiSupCon was the only strategy that led to a reasonable accuracy.

The results of this study indicate that training a model using a self-supervised approach leads to low predictive performance, while fully supervised training results in poorly trained models due to limited availability of annotated data. In contrast, our semi-supervised approach achieved higher predictive performance by leveraging a larger pool of unannotated images and incorporating bioactivity annotations during training.

The importance of providing neural networks with ground truth annotations for representation learning is further underscored when comparing the SemiSupCon results on the BBBC022 and BBBC036 datasets. Despite the BBBC036 dataset containing an order of magnitude more images than BBBC022, the downstream accuracy on the BBBC036 dataset is substantially lower for all evaluated tasks compared to downstream classification on the BBBC022 dataset. This discrepancy could be explained by the substantially lower annotation ratio of MeSH classes in the BBBC036 dataset compared to BBBC022. The low annotation ratio results in the self-supervised contrastive loss having a substantially greater impact on the model’s weight updates than the supervised contrastive loss.

Since SemiSupCon also used MeSH classes to define positives during the representation learning phase of the workflow, it is within expectations that predictive performance on annotations from the Drug Repurposing Hub would be lower. SemiSupCon also demonstrated considerable predictive performance on new annotations, indicating that the model is capable of learning phenotypical representations from the microscopy images that also can be applied to new bioactivity prediction tasks.

SemiSupCon performed worse at predicting Drug Repurposing Hub target labels compared to the Drug Repurposing Hub MoA labels, as particularly evident in the multi-label accuracies. Not only was the accuracy for predicting target labels low, but it also exhibited a high standard deviation. The poorer performance in predicting target labels likely stems from the higher number of different target classes compared to MoA classes (see Table S1). A greater number of possible classes particularly impacts the multi-label accuracy, as all classes for a given data point have to be correctly predicted.

To further examine the robustness and adaptability of our workflow, we also evaluated an inductive SemiSupCon strategy, which slightly improved downstream classification of MeSH classes and Drug Repurposing Hub MoA annotations (Table S2). This showcases that SemiSupCon is capable of learning discriminative phenotypical representations and classifiers trained with these representations can potentially be used to predict different annotation systems.

While our current study primarily uses MeSH classes as labels during the training of SemiSupCon, these results suggest a potential strategy to enhance the performance of our workflow, particularly on the BBBC036 dataset. This strategy could involve incorporating additional biological annotations of compounds during the training of SemiSupCon to reduce the influence of the self-supervised contrastive loss. Annotations from the Drug Repurposing Hub could be included into the training process, or additional annotations could be extracted from other public databases, such as ChEMBL^29^ or Probes & Drugs^93,94^.

The predictions of our workflow were deconvolved for each MeSH class to identify for which specific classes our workflow demonstrated robust predictive performance. Our results showed that our workflow could predict 106 out of 253 MeSH classes in BBBC022 dataset with sufficient precision (10 % was chosen as threshold), suggesting a wide range of bioactivity could be inferred from image-based profiling data. Several potential reasons may explain why our workflow failed to accurately predict certain MeSH classes, while high precision could be achieved for others. One possibility is that bioactive compounds from some pharmacological classes do not induce significant morphological perturbations in cells, while there are pharmacological classes which can induce strong morphological perturbations.^95,96^ Additionally, the U2OS cells used in these two datasets may not be the most suitable model system for every pharmacological class. Among the top 25 MeSH classes with the highest predictive performance, we identified several associated with biological activities unrelated to human cells. This aligns with a recent study that predicted compound bioactivity from Cell Painting data on human cell lines, in which several bioactivities were linked to yeast and bacterial targets.^97^ The non-human-cell-related MeSH classes identified in our study include anti-bacterial agents, anti-fungal agents, herbicides and insecticides. Given the polypharmacology of many compounds, it is plausible that these small molecules have targets within human cell lines. For example, several hypoglycemic agents are organic compounds in the sulfonylurea family (e.g. Chlorpropamide^98^), known for their use as herbicides.^54^ While 2,4,5-trichlorophenoxyacetic acid, an herbicide, has no direct evidence for being a hypoglycemic agent, its close derivative 2,4-dichlorophenoxyacetic acid is an herbicide and a hypoglycemic agent^55^. Indeed, when visualizing the image representation of annotated CP dataset via PHATE, it is evident that herbicides and hypoglycemic agents form clusters that are closely positioned to each other in the phenotypical space (Figure 4C).

We also explored the learned phenotypical space independent from the annotations of the compounds, to identify regions where compounds with similar pharmacological activities were clustering together. By doing this, we identified a cardiac glycoside and a tubulin modulator cluster. The two clusters were also identified by Gustafsdottir *et al*.^42^, but it is worth noting that using our approach we were able to cluster more tubulin modulators and cardiac glycosides together in their respective cluster.

Our workflow demonstrated its potential in classifying the biological activity of annotated compounds. We then proceeded to use the downstream classifiers to predict single-label MeSH classes of unannotated compounds, which were ranked based on the batch-wise calculated contrastive loss. In the BBBC022 dataset, we successfully confirmed predictions for 7 out of 10 compounds, where the predictions from the replicates exhibited overall high agreement (Figure 5). Multiple MeSH classes were predicted for these compounds, and for most of them evidence was found linking them to multiple of the predicted MeSH classes. To better assess the reliability of our predictions in the future, more advanced deep learning techniques such as Monte Carlo dropout^99^ could be used to approximate the uncertainty of the predictions.

Although the downstream accuracy of SemiSupCon(BBBC036) was much lower compared to the same workflow trained on the BBBC022 dataset, we were still able to correctly predict MoA for unannotated compounds in the BBBC036 dataset, which were verified through a literature search (Figure 6). The low accuracy reflects the limited ability of this workflow to accurately predict a wide range of MeSH classes, but a few MeSH classes can nonetheless be accurately predicted. This is highlighted by the fact that 4 from 6 of validated compounds belong to the top 7 most predictable MeSH classes for this workflow.

## 5 Conclusion

In this study, we introduced a semi-supervised contrastive learning method to learn representations of the morphological perturbations induced by small molecules in a Cell Painting assay. We utilized image replicates as positive pairs during self-supervised contrastive learning and additionally the MeSH classes of annotated compounds in subsequent supervised contrastive learning. In downstream classification tasks, we employed the learned image representations to evaluate MeSH class predictions, as well as to predict MoAs and targets from the Drug Repurposing Hub. We demonstrated that SemiSupCon improved downstream classification compared to Con, DINO and expert-engineered CellProfiler profiles, showcasing the utility of transductive semisupervised learning for bioactivity prediction. Using this workflow, we identified that over 106 MeSH classes could be predicted with sufficient precision on the annotated dataset of BBBC022.

Notably, this workflow enabled us to predict the bioactivity of a number of unannotated compounds, some of which were later confirmed through literature searches. Our workflow offers a practical strategy for using Cell Painting images for bioactivity prediction with minimal human intervention.

Previous studies have already demonstrated that incorporating features from additional data modalities can increase predictive performance, an approach we also plan to include in our future works.^11,97^ Especially, graph neural networks stand out as powerful and versatile tools for learning suitable representations, and they have proven to be highly effective in learning unified representations^100,101^ by fusing knowledge from multiple modalities (e.g. gene expression^102,103^ or molecular structure^104,105^). Additionally, the proposed semi-supervised model could be further enhanced by integrating bioactivity information from various annotation systems. With these enhancements, semi-supervised contrastive models trained on CP data could become valuable tools for routine use in *in silico* phenotypical screening, identifying compounds with adverse side effects, drug repurposing, and uncovering the bioactivity of natural products.

## Data and code availability

The data and code to reproduce our results are available at our GitHub repository (https://github.com/AGSun-FMP/CP_SemiSupCon).

## Author Contributions

D.B., C.S. and H.S. designed and directed the project with contributions from all authors. D.B. constructed the machine-learning models and analyzed the data. C.B. generated the CellProfiler profiles. D.B. and H.S. wrote the manuscript with contributions from all authors.

## Acknowledgements

This work was supported by the Leibniz-Forschungsinstitut für Molekulare Pharmakologie (FMP), and the Deutsche Forschungsgemeinschaft (DFG, German Research Foundation) under Germany’s Excellence Strategy–EXC 2008/1 (UniSysCat; 390540038) (to H.S.). We thank Dr. J. P. von Kries for helpful discussions.

