## Supplemental Information for "Semi-supervised Contrastive Learning for Bioactivity Prediction using Cell Painting Image Data"

Table S1. Total number of annotations and number of unique annotations for MeSH class and repurposing hub annotations.

| Annotations | BBBC022 | BBBC036 |
| --- | --- | --- |
| Compounds with MeSH classes | 839 | 1079 |
| Unique MeSH classes | 298 | 317 |
| Unique single-label MeSH classes | 223 | 253 |
| Compounds with Drug Repurposing Hub MoA annotations | 773 | 1534 |
| Unique Drug Repurposing Hub MoA classes | 291 | 592 |
| Compounds with Drug Repurposing Hub target annotations | 587 | 720 |
| Unique Drug Repurposing Hub target classes | 659 | 774 |

**Table S2.** An inductive SemiSupCon approach was compared against baselines on downstream classification on the BBBC022 dataset. CellProfiler, transductive Con, transductive DINO and a fully supervised trained ResNet50 were used as baselines. Downstream MLP was evaluated for multi-label prediction of MeSH classes and Drug Repurposing Hub annotations for all methods except for the supervised trained ResNet50. The supervised ResNet50 was only trained and evaluated for multi-label MeSH class classification (5-CV used). 80 % of the compounds were in the training set and the remaining 20 % were in the test set.

| Strategy | MeSH class accuracy [%] | Drug Repurposing Hub MoA accuracy [%] | Drug Repurposing Hub targets accuracy [%] | |
| --- | --- | --- | --- | --- |
| CellProfiler | 0 | 0 | | 0 |
| DINO-tiff | 0 | 0 | | 0 |
| Con | 0.39 | 0.59 | | **0.64** |
| SemiSupCon-inductive | **0.64** | **0.86** | | 0 |
| Supervised ResNet50 | 0 | **-** | | - |

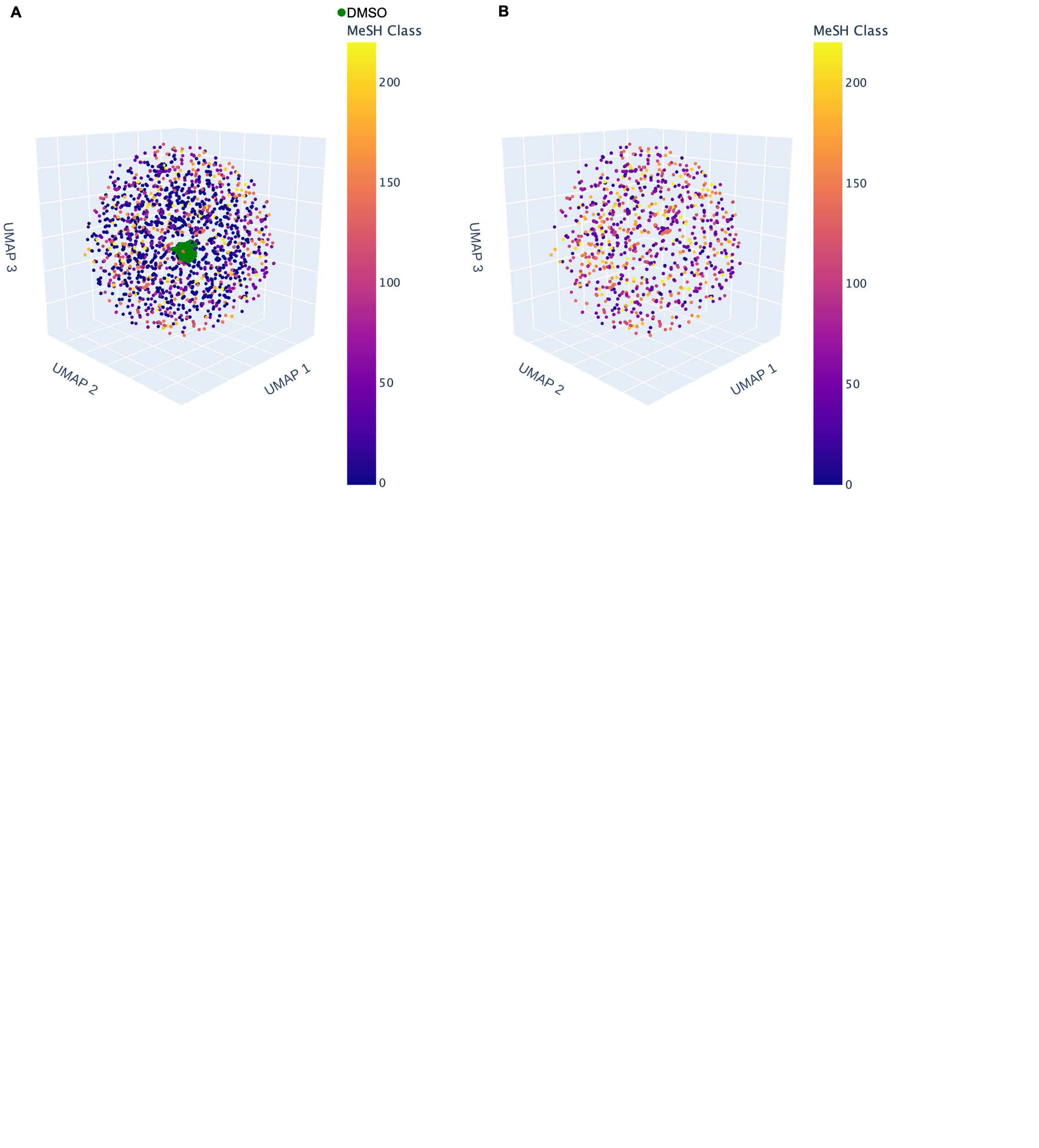

Figure S1. 3D visualization of the phenotypical space from the SemiSupCon(BBBC022) model with UMAP. Data points were colored according to their MeSH classes (compounds with multiple MeSH classes were assigned to their first MeSH class). Lowest color code indicates unavailability of a MeSH class, while green data points represent the DMSO controls. (A) Visualization with all data points. (B) Visualization with only the annotated subset.

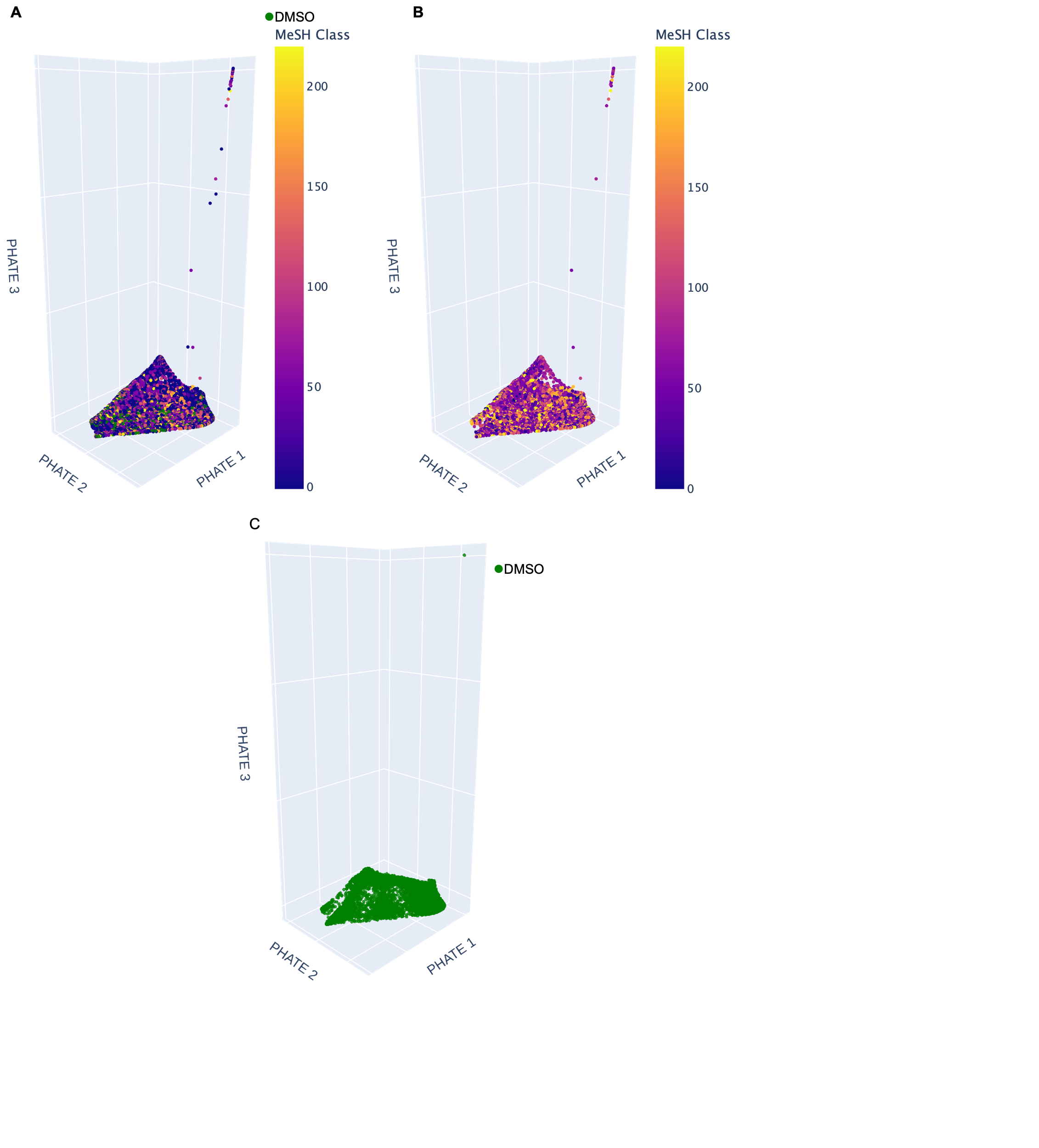

**Figure S2.** PHATE visualization of the CellProfiler profiles calculated from the BBBC022 dataset. Data points are colored according to their MeSH classes (compounds with multiple MeSH classes were assigned to their first MeSH class). Lowest color code indicates unavailability of a MeSH class, while green data points represent the DMSO controls. (A) Visualization with all data points. (B) Visualization with only the annotated subset. (C) Visualization with only the DMSO control.

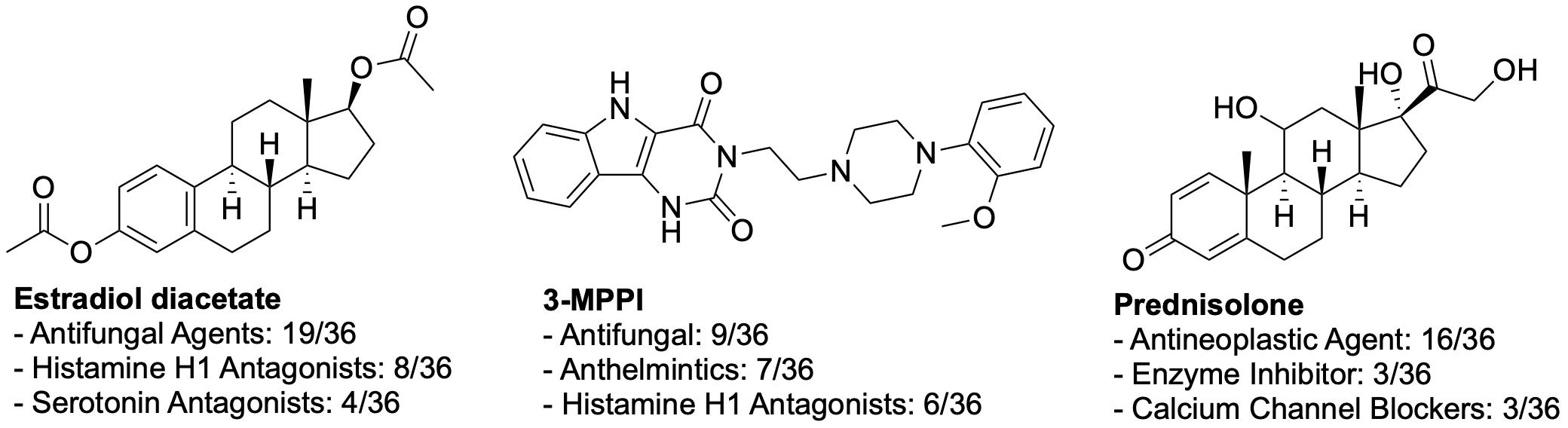

**Figure S3.** Compounds from top 10 unannotated compounds in the BBBC022 dataset with the lowest contrastive loss, where predictions of MoAs could not be validated through a literature search using a single-label RF model. Predictions were made for each replicate of a compound (36 in total), and for each compound, the three MeSH classes with the highest number of predictions are shown.

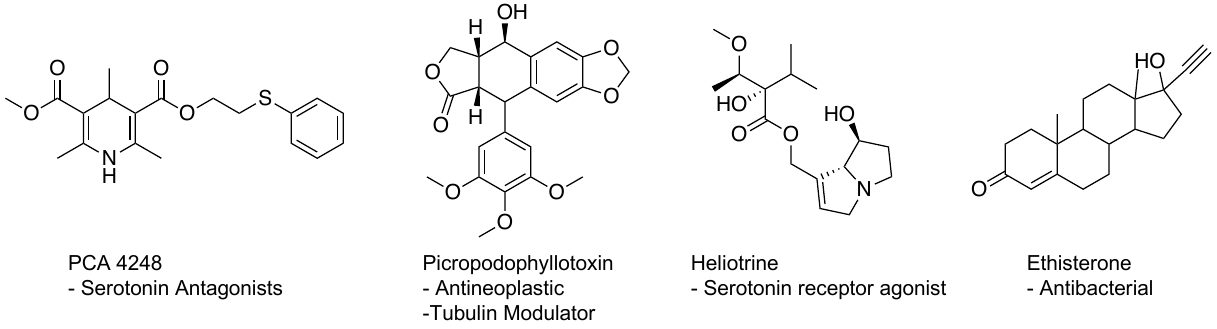

**Figure S4.** Four predictions of MoAs, confirmed through literature search, for the 10 unannotated compounds with the lowest contrastive loss in the BBBC022 dataset using a multi-label RF model trained on SemiSupCon(BBBC022) representations. PCA 4248 was identified to be a serotonin antagonist^1^. Picropodophyllotoxin was confirmed to be both a tubulin modulator and an antineoplastic compound^2^. Heliotrine was confirmed to be a serotonin receptor agonist^3^ and Ethisterone was confirmed to have antibacterial activity^4^.

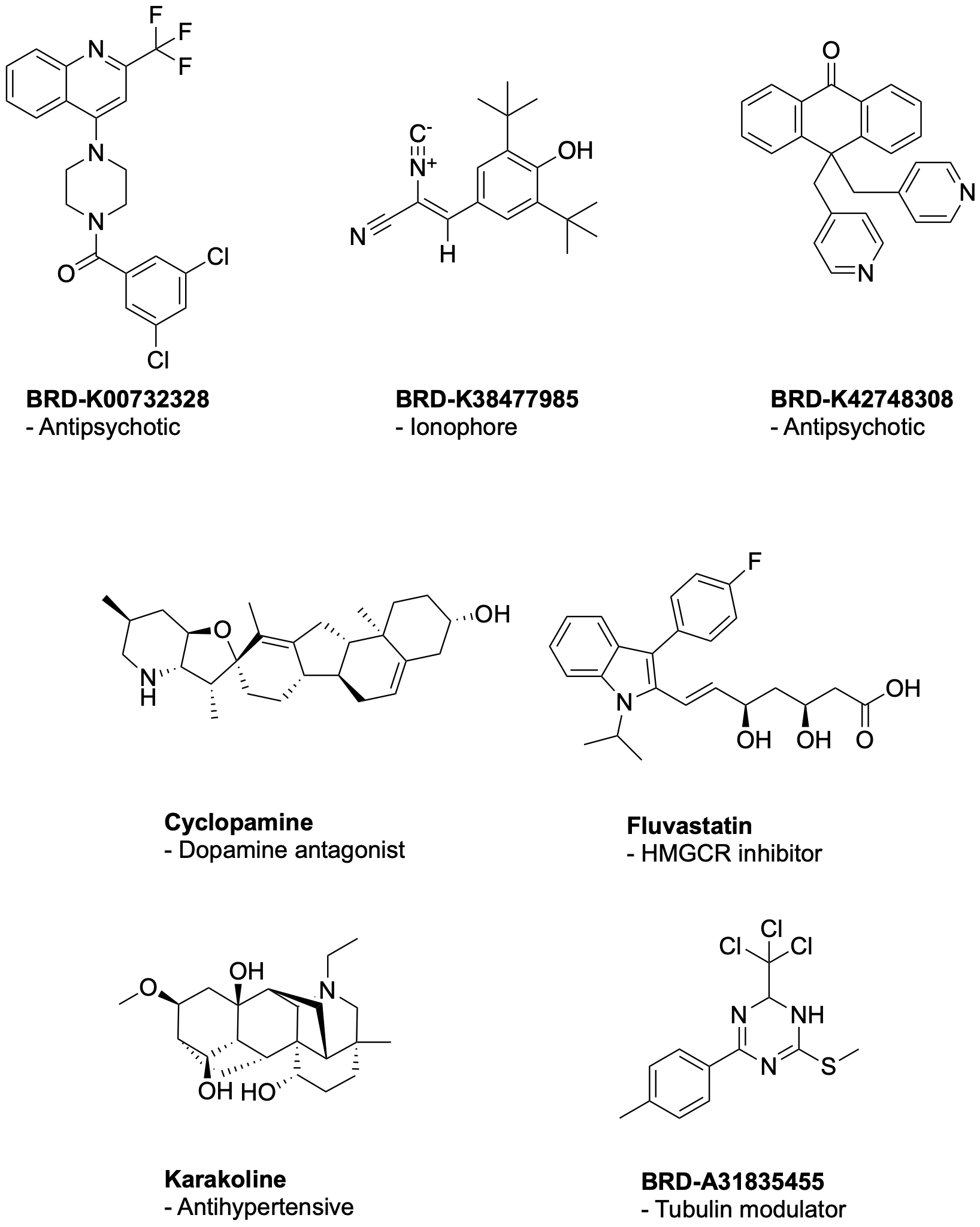

**Figure S5.** Seven MoA predictions that were confirmed through literature searches from top 25 profiles of unannotated compounds in the BBBC036 dataset with the lowest contrastive loss. The predictions were generated using a multi-label RF model trained on SemiSupCon(BBBC036) features. Specifically, BRD-K00732328 was confirmed as an antipsychotic compound^5^, BRD-K38477985 as an Ionophore^6^, BRD-K42748308 as an antipsychotic compound^7^, Cyclopamine as a dopamine antagonist (Cyclopamine inhibits the hedgehog pathway)^8^, Fluvastatin as a HMGCR inhibitor^9^. Evidence suggests Karakoline functions as an antihypertensive compound^10^, and tubulin as one of the targets of BRD-A31835455 in ChEMBL^11^.
